# Combined Transplantation of Mesenchymal Progenitor and Neural Stem Cells to repair cervical spinal cord injury

**DOI:** 10.1101/2025.02.13.638141

**Authors:** Seok Voon White, Yee Hang Ethan Ma, Christine D Plant, Alan R Harvey, Giles W Plant

## Abstract

Mesenchymal progenitor cells (MPC) are effective in reducing tissue loss, preserving white matter and improving forelimb function after spinal cord injury (SCI) (White et al., 2016). We proposed that by preconditioning the mouse by intravenous delivery (IV) of MPCs for 24 hours following SCI that this would provide a more favorable tissue milieu for NSC intraspinal bridging transplantation at day 3 and day 7. In com-bination these transplants will provide better anatomical and functional outcomes. The intravenous MSCs would provide cell protection and reduce inflammation. NSCs would provide a tissue bridge for axonal regeneration and myelination and reconnect long tract spinal pathways. Results showed that initial protection of the injury site by IV MPCs transplantation resulted in no increased survival of the NSCs transplanted at Day 7. However, integration of transplanted NSCs was increased at the Day 3 time point indicating MPCs influence very early immune signaling. We show in this study that MPC transplantation resulted in a co-operative NSC cell survival improvement at Day 3 post SCI. In addition to increased NSC survival at day 3 there was an increase in NSC derived mature oligodendrocytes at this early time point. In vitro analysis confirmed MPC driven oligodendrocyte differentiation which was statistically increased when compared to control NSC only cultures. These observations provide important information about the combination, delivery, and timing of two cellular therapies in treating SCI. This study provides important new data on understanding the MPC inflammatory signaling within the host tissue and time points for cellular transplantation survival and oligodendroglia differentiation. These results demonstrate that MPC transplantation can alter the therapeutic window for intraspinal transplantation by controlling both the circulating inflammatory response and local tissue milieu.

## 1. Introduction

Spinal cord injury (SCI) pathophysiology is extensive and can include cell death, neuro-inflammation, oxidative stress, demyelination, and axonal degeneration. Given these multiple outcomes post injury, a combinatorial approach to SCI treatment would provide an excellent way to target the numerous positive changes required for recovery of function. Previous combinational strategies have included cellular transplantation mixed with drugs and/or growth factor treatments. Adult cellular transplantation strategies have included Schwann cells, olfactory ensheathing glia, macrophages, neural stem cells (NSCs), induced pluripotent stem cells and mesenchymal progenitor cells (MPCs) [1–4]. MPCs or more commonly used mesenchymal stem cells (MSCs) have been studied extensively as a therapeutic agent for SCI, second only to Schwann cells [2]. Among the benefits of using MPCs is that they are immunomodulatory and neuroprotective to the injured spinal cord [5, 6]. However, there is a limited number of reports that show MPCs can promote regeneration of injured axons, act as a long-term tissue bridge or relay grafts, and promote synaptic reconnections [7]. For this reason, additional bridging strategies will be required to bridge the tissue gap left by SCI.

NSCs have been identified in the adult brain and spinal cord, successfully cultured in vitro and can differentiate into neurons, astrocytes or oligodendrocytes [8]. This has allowed further study of these cells not only endogenously, but also following their transplantation in the experimental setting of SCI [9–14]. Transplantation of NSCs to the cervical spinal cord have resulted in some functional improvements and survival of up to 10 weeks, but the usual default differentiation in vivo are astrocytes [9]. Neural progenitor transplantation has also been shown to be beneficial in treating SCI when used in combination with neurotrophic factors [15]. These studies indicate that NSCs have significant repair and regenerative potential when used in SCI.

A major factor in the viability and phenotype of transplanted cells following transplantation to the spinal cord is the inflammatory milieu after injury. Due to the acute inflammatory cascade, many intraspinal cell injections take place seven to fourteen days after injury [16, 17]. We have recently shown that intravenous injection (IV) of MPCs 24 hours following injury decreases tissue loss, angiogenesis, glial and pericyte scarring and cellular inflammation after SCI [18]. No evidence of any increased axonal regeneration was achieved by IV delivery of MPCs [19] and no MPCs were found to have integrated into the spinal cord at any segment level. This indicates a therapeutic need to re-connect the rostral and caudal aspects of the lesion site by a cellular or material bridge in conjunction with the therapeutic MPC treatment. For that reason we have explored the co-transplantation of MPCs (Intravenously) with NSCs (intraspinally) to provide an optimal regenerative combination. Dual transplantation techniques have been developed include adipose-derived mesenchymal stem cells with NSCs which promoted greater survivability of NSCs after transplantation [20].

In this study we have investigated a combinatorial cellular transplantation approach utilizing an early intervention (Day 1) IV injection of Sca-1+ selected MPCs [21] with intraspinal injection of NSCs (Day 3 and Day 7) into a cervical SCI model. We hypothesized that this cellular combination would utilize the anti-inflammatory capacity of MPCs and tissue bridging capacity of NSCs to: (1) protect the injured cervical spinal cord by reducing tissue loss and immune cell infiltration; (2) provide a bridging/relay graft for regenerating axons and (3) reduce innate inflammation, thereby influencing transplanted cell differentiation. We also hypothesize that IV injection of MPCs at 24 hours post SCI can provide an earlier intervention window for intraspinal injections based on our previous observations [19]. Intraspinal NSCs injections at D3 and D7 post-injury were chosen to provide transplant days with high and lower levels of inflammatory cell responses [22]. Overall, we postulate that IV MPC delivery should significantly improve tissue milieu for NSCs survival, integration and provide a synergistic therapy for return of forelimb function.

## 2. Materials and Methods

### Animals

Female FVB mice (12-14 weeks, Charles River) were used for all surgeries and cell injections. For primary MPC cultures, adult FVB mice were used. Primary NSC cultures were obtained from adult transgenic mice ubiquitously expressing green fluorescence protein (GFP) and firefly luciferase reporter gene (lucGFP) driven by a chicken β-actin promoter [23, 24] with an FVB background (gift from Professor Joseph Wu, Stanford, USA). All animals were housed in a clean barrier facility on a 12/12-h dark-light cycle. Stanford University Administration Panel approved all protocols on Laboratory Animal Care (APLAC) committee per IACUC guidelines. The experimental design is as shown in Figure 1A.

**Figure 1.**
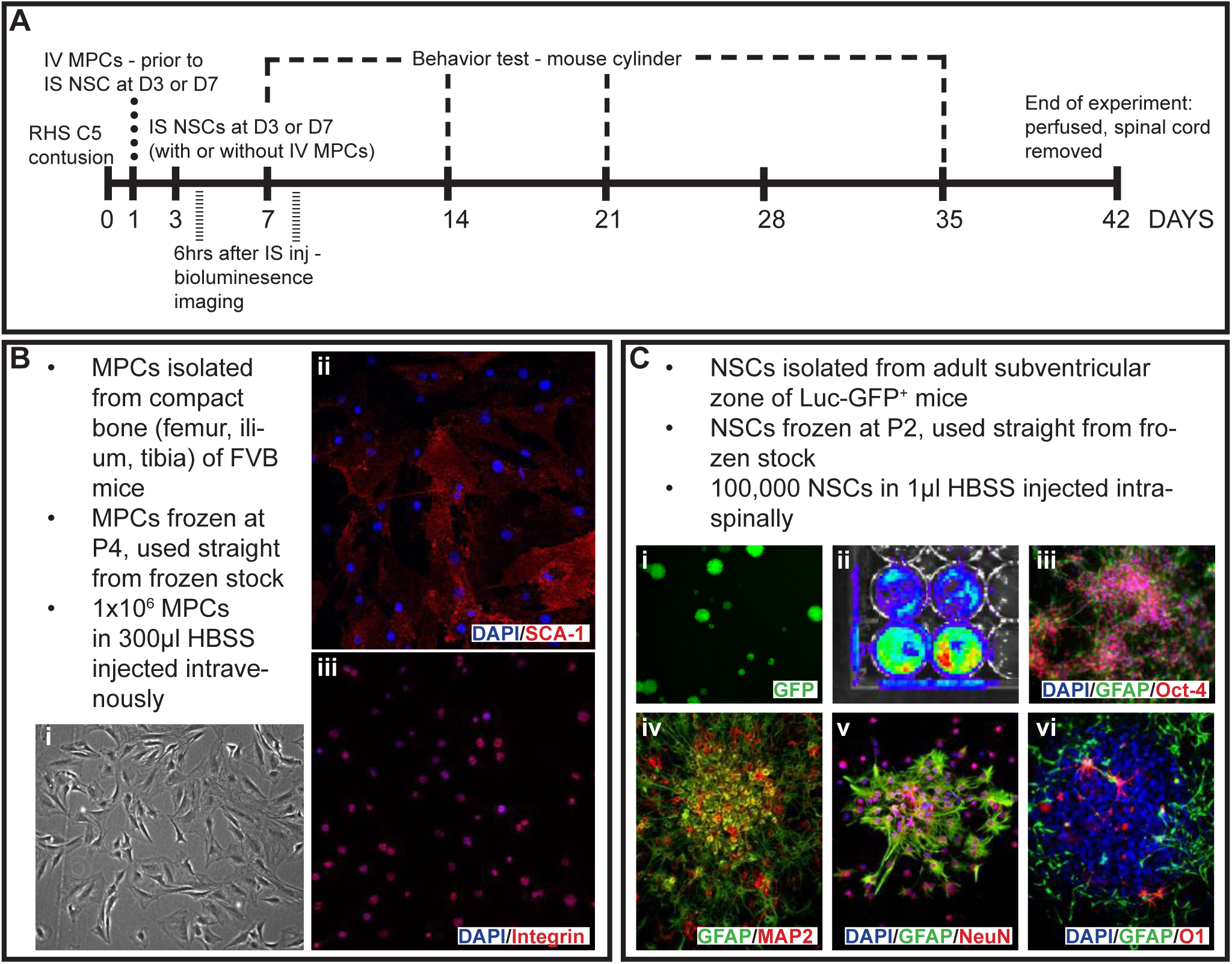
Schematic of experimental design and primary cell cultures used in the study. (A) Outline of the experimental design, timing of injections and key time points in the study. Cells were obtained from primary cell cultures including (B) MPCs isolated from the compact bone of FVB mice with (i) phase image of MPCs in culture. Isolated MPCs were positive for (ii) Sca-1 and (iii) integrin. (C) Primary cell cultures of NSCs were isolated from the adult subventricular zone of Luc-GFP+ mice with (i) showing endogenous expression of GFP, (ii) bioluminescence signal, spontaneous differentiation of adhesive NSCs with (iii) GFAP and Oct-4 and (iv) GFAP and MAP-2. NSCs when co-cultured in a non-contact mediated manner with MPCs differentiated into cells expressing (v) NeuN and GFAP and (vi) GFAP and O1 for neurons, astrocytes and oligodendrocyte identification. Scale bars = 20 µm.

### Mesenchymal Progenitor Cell Culture

MPCs were isolated from the compact bone of female FVB mice as adapted from [21] and previously used in White et al., 2016 [19]. Briefly, 10 adult mice were euthanized with an overdose of sodium pentobarbital (Beuthanasia-D, 0.01ml/30g). The ilium, femur and tibia were removed, then cleaned and crushed to release the cells; blood or bone marrow cells were discarded. The resulting cell suspension was depleted of CD5, CD45R, CD11b, Anti-Gr-1, 7-4, Ter-119 and CD3ε positive cells on an autoMACs Pro Separator (Miltenyi Biotech, USA) using the PosselD2 program. The resulting negative cell population was then selected for Sca-1 (Miltenyi Biotech, USA) with an autoMACs Pro Separator using the PosselD2 program. Sca-1+ MPCs were plated down in media consisting of αMEM, 20% fetal calf serum, 1x GlutaMAXTM, 1x sodium pyruvate 100mM and 1mg/mL gentamicin solution (all reagents from Life Technologies, USA) at a density of 10000 cells/cm2 and expanded until P4 (Figure 1B). Cells were frozen at 1x106 cells/ml of media with 7% DMSO and stored in liquid nitrogen until required.

### Neural Stem Cell Culture

NSCs were isolated from the subventricular zone of female lucGFP mice as adapted from Azari and colleagues [25]. Briefly, 5 adult mice were euthanized with a sodium pentobarbital (Beuthanasia-D, 0.01ml/30g) overdose. The brain was removed and the thin layer of tissue surrounding the lateral wall of the ventricles was cut, excluding the striatal parenchyma and corpus callosum. The resulting tissue was minced and enzymatically digested with 3ml of 0.05% trypsin-EDTA (Life Technologies, USA) for 7 min at 37°C. The resulting cell suspension was mechanically dissociated. Cells were washed, passed through a 40 µm cell strainer, then plated down into one T25 flask per brain in 5ml of complete NSC medium supplemented with 20 ng/ml epidermal growth factor, 10 ng/ml basic fibroblast growth factor and 1µl/ml of 0.2% heparin (all items from StemCell Technologies, Canada) and expanded until P3 as free-floating neurospheres. Cells were then frozen as per guidelines from StemCell Technologies. One flask was frozen per tube in 1.5 ml of media with 10% DMSO.

### Surgeries

An average of 10 mice was used per group. The injury model used was a contusion injury at cervical level 5 using an Infinite Horizon impactor (Precision Systems and Instrumentation, USA). Mice were deeply anesthetized using isofluorane (2.5% in O2) and a C5 laminectomy was performed to expose the spinal cord. A custom-made 1mm impactor head was used to deliver the impact. C6 and C4 vertebra were clamped. The impactor head was aligned to the midline of the exposed spinal cord and moved 2mm (one rotation) to the right. An impact of 30kDy with 3s dwell was delivered. The muscles and skin were sutured. Postoperative care consisted of the administration of Pfizerpen (penicillin G potassium, 250,000units/mL, SC, Novaplus, Cat#: NDC0049-0520-22), buprenorphine (0.01mg/kg SC, twice a day for 3 days) and saline (1ml/20g, twice a day for 3 days).

### MPCs Preparation for IV Injections

Previously frozen MPCs were rapidly thawed (37°C water bath) and transferred to 15mL tubes containing 5mL of Hank’s Balanced Salt Solution (HBSS; Life Technologies, USA). Tubes were centrifuged for 5 mins at 400g and the resulting pellet resuspended into a single-cell suspension with HBSS. The wash step was repeated to remove excess DMSO. Cell pellet was resuspended in 300µL of HBSS and transferred to a 1.5mL microcentrifuge tube and kept at room temperature prior to IV injections. Cell viability was checked using trypan blue with random samples taken before and after IV injections. Cells were >99% viable pre-injection and were >85% viable post-injection.

### Intravenous injections of MPCs

In the cohort of mice receiving dual injections, MPCs were injected intravenously at D1 post-injury via the tail vein. Mice were lightly anesthetized using isofluorane (1.5% in O2). The tail was cleaned with alcohol and heated until the vein was dilated. MPCs were resuspended into a single cell suspension and transferred to 1mL syringe. MPCs were injected into the tail vein using a 30g needle. For control mice, 300µL of HBSS was injected into the tail vein.

### NSCs Preparation for intraspinal injections

Frozen NSCs were rapidly thawed (37°C water bath) and transferred to 15 ml tubes containing 5 ml of Hank’s Buffered Saline Solution. Tubes were centrifuged for 5 mins at 110 x g and the resulting pellet resuspended in HBSS into a single cell suspension. The wash step was repeated to remove excess DMSO. The resulting cell suspension was counted and resuspended in appropriate amounts of HBSS to ensure 100,000 cells/µl of HBSS. Cell viability was checked using trypan blue before and after injections. Cells were >98% viable pre-injection, and >80% viable post-injection.

### Intraspinal Injections of NSCs

Animals receiving intraspinal injections at D3 or D7 post-injury were anesthetized using isofluorane (2.5% in O2) and the spinal cord was exposed at the previous injury site. Vertebra level C6 and C4 were clamped to straighten the spinal cord for injection. A Nanoject IITM (Drummond Scientific, USA) with a custom glass pipette tip was used to inject 100,000 cells in 1 µl of HBSS at 200nl/min into 0.8 mm lesion epicenter. For control mice, 1 µl of HBSS was injected at 200nl/min into 0.8 mm lesion epicenter. After injection, the needle was left in place for 2 mins before retracting. For control animals, 1 µl of HBSS was injected at 200nl/min into 0.8 mm lesion epicenter. The muscles and skin were sutured. Postoperative care consisted of the administration of Pfizerpen (penicillin G potassium, 250,000units/mL, SC), buprenorphine (0.01mg/kg SC, twice a day for 2 days) and saline (1ml/20g, twice a day for 3 days).

### Group designation

After histological processing, mice that had injuries crossing over the midline were omitted from the overall results reported. Supplementary Table 1 shows the abbreviations used in text for each group, as well as the total number of animals used per group for analysis (n).

### Bioluminescence

Mice receiving NSC injections were imaged at 6 hrs post-injection for a bioluminescence signal using the IVIS Spectrum (PerkinElmer, USA), to track the injected NSCs. Mice were lightly anesthetized using isofluorane (1.5% in O2) and injected with D-luciferin K+ Salt Bioluminescent Substrate (150mg/kg; PerkinElmer, USA). Five minutes after injection, a series of four 2-minute exposures and one 3-minute exposure with medium binning at F/stop 1 was taken. Bioluminescence emissions were calculated using Living Image software (V4.3.1) ROI contour tool with threshold at 20%.

### Behavior - Mouse cylinder

Behavioral data was collected using the mouse cylinder test for gross paw usage [26]. All mice were recorded at D7, D14, D21 and D35. Mice were placed in a Perspex cylinder and the number of left and right paw touches were recorded for 5 mins; only full touches with hind leg rearing were counted as a true result for analysis. The percentage of left versus right paw usage was then calculated (right-paw usage/left + right-paw usage x 100%).

### NPCs and MSCs culture

NPCs and MSCs are first grown in separate cultures. NPCs are isolated from the hippocampal tissues of female adult (10-12weeks) C57BL/6 mouse (gift of Theo Palmer’s lab) and frozen in nitrogen until use. NPCs are rapidly thawed (37oC) and grown in mNPC media consisting of Neurobasal-A medium supplemented with 1X B27 without Vitamin A (Invitrogen), 20ng/ml bFGF (Peprotech), 20ng/ml EGF (Peprotech), and 1X GlutaMAX (Gibco). NPCs are grown on poly-l-lysine (10ug/ml; Sigma-Aldrich) and laminin (5ug/mL; Sigma-Aldrich) coated 10cm dished. NPCs are split every 3-4 days using 0.25% Trypsin + 0.5mM EDTA and plated at a density of 1.0x105 cells per dish. MSCs are isolated from compact bone of adult female FVB mice (12-14 weeks, Charles River) and grown in media consisting of 10% HyClone fetal Bovine serum, 1X GlutaMAX, and 1mM Sodium Pyruvate (Gibco).MSCs were fed with fresh media every 3 days and split at 60% confluency using 0.05% Trypsin +0.5mM EDTA.MSCs are expanded using low plating density in Corning T75 tissue culture flasks until p4 and subcultured into Corning 24-well Transwell inserts for co-culture.

### MSC driven NSC differentiation

NSCs are seeded onto poly-l-lysine (10ug/ml; Sigma-Aldrich) and laminin (25ug/mL; Sigma-Aldrich) coated glass coverslips at a density of 120,000 cells/well and allowed to grow overnight with complete NPC growth media. P4 MSCs are split and plated onto Corning 24-well transwell inserts containing a porous polyester membrane (0.4um pores) in MPC growth media overnight. The next day, NPC media is replaced with NPC differentiation media consisting of Neurobasal-A medium supplemented with B27w/Vitamin A (Invitrogen), 1X GlutaMAX, and 0.5ng/ml EGF (Peprotech), 1mM sodium pyruvate (Gibco), and 1X MEM non-essential amino acids (Gibco). Transwell MSCs are inserted to wells containing NPC glass coverslips for 6 hours, 12 hours, 24 hours, 7 days, or 10 days. Half-media changes of fresh NPC differentiation media and MPC 10% FBS growth media were performed every 2 days before immunocytochemical analysis on 10th day of differentiation. NPC differentiation media or NPC differentiation media plus Transwell insert containing the MPC complete media with no cells were done as control.

### Immunocytochemistry

Following 10 days of differentiation, NPC identity was analyzed via immunocytochemistry staining. Differentiated NPC coated glass coverslips were washed three times with 1x PBS and incubated in 0.1% Triton C-100 in PBS for 10 minutes at room temperature. After, coverslips were washed with PBS and incubated in blocking buffer (3% normal donkey serum in PBS) for one hour. Following blocking buffer, coverslips are incubated with designated primary antibodies for 4 hours at room temperature (Table 1). Coverslips were washed with 1x PBS and subsequently incubated with the corresponding secondary antibodies for each species for 2 hours. Secondary antibodies are obtained from Jackson ImmunoResearch Laboratories and consisted of Alexa Fluor 488 (1:250), Cy 3 (1:250), or Cy 5 (1:250) diluted in blocking buffer solution containing DAPI. Glass coverslips are then washed with 1x PBS and mounted using Fluormount G (Southern Biotech) and imaged using Nikon C2 confocal microscope.

**Table 1.** Primary Antibodies.

| Primary Antibody | Supplier | Host | Dilution |
| --- | --- | --- | --- |
| Beta-III Tubulin | Aves Lab, Inc | Chicken | 1:1000 |
| O4 | EMD Millipore | Mouse | 1:50 |
| GFAP | Dako | Rabbit | 1:500 |
| MAP2 | Aves Lab, Inc | Chicken | 1:1000 |
| APC | CalBiochem | Mouse | 1:200 |
| GFAP | Dako | Rabbit | 1:500 |
| TuJ1 | Biolegends | Mouse | 1:500 |
| NG2 Chondroitin Sulfate | EMD Millipore | Rabbit | 1:500 |
| GFAP | Aves Lab, Inc | Chicken | 1:500 |

### Histology

At D42, animals were euthanized with an overdose of Beuthanasia-D and transcardially perfused with 4% paraformaldehyde. Spinal cord tissue was collected and post-fixed in 4% paraformaldehyde overnight, then switched to 30% sucrose in PBS. Cords were sectioned horizontally on a freezing microtome (Leica, USA) at 50µm thickness.

### Immunostaining (Tissues)

Tissue sections were blocked for 1h at room temperature in PBS + 10% normal donkey serum + 0.2% Triton X-100 (diluent). Sections were incubated in primary antibody in diluent for 48 hours at 4°C and secondary antibody in diluent for 30 minutes at room temperature. Primary antibodies used were βIII tubulin (mouse, 1:800; Covance, USA), glial fibrillary acidic protein (GFAP, rabbit, 1:500; Dako, USA), adenomatous polyposis coli (APC; mouse, 1:200; Abcam, USA), chondroitin sulfate (CS-56, 1:800, Sigma-Aldrich, USA), beta-type platelet-derived growth factor receptor (PDGFrβ, rabbit, 1:200, Abcam), p75 neurotrophin receptor (p75, rabbit, 1:400, Promega, USA) and green fluorescence protein (GFP, chicken, 1:10000; Aves Lab Inc, USA). The corresponding secondary antibody for each species was from Jackson Laboratories and included AF647 anti-mouse (1:800), DyLightTM 549 anti-rabbit (1:1000), and DyLightTM 488 anti-chicken (1:800). Blood vessels were labeled with DyLight® 488 Lycopersion Esculentum (Tomato) Lectin (1:100; Vector Laboratories, USA). Myelin was labeled with FluoroMyelin (1:300, Life Technologies, USA). Fluorescence images were taken using a Nikon C2 confocal microscope.

### Intraspinal injection NSC counts

Using sections labeled with anti-βIII tubulin, anti-GFAP and anti-APC and cellular morphology, GFP+ cells were counted in 8 sections per mouse using 2 blinded observers. The counted cells were categorized as neurons, astrocytes or oligodendrocytes based on the antibodies described above.

### Statistics

#### In vivo analysis

One-way ANOVA with Tukey’s post-hoc test statistics were performed using GraphPad Prism Software. Values of P < 0.05 were considered statistically significant. All error bars were standard deviation.

#### In vitro Analysis

All statistical analyses were conducted using the RStudio software (2025). Data were presented with error bars indicating Standard Error of Mean (SEM). Depending on data characteristics, either Welch’s one-way analysis of variance (Welch’s ANOVA) or standard ANOVA was used for F-tests. Post hoc comparisons were performed using Tukey’s HSD, or Dunn’s Test, as appropriate. p ≤ 0.05 denotes statistical significance. Significance was also presented with the following notations: * (p ≤0.05), ** (p ≤0.01), *** (p ≤0.001), **** (p ≤0.0001).

## 3. Results

### Bioluminescence imaging is an effective tracker of intraspinally injected NSCs

Luc-GFP+ NSCs were injected intraspinally at D3 or D7 post-injury with or without IV injection of MPCs at D1 post-injury; cell fate was tracked using bioluminescence imaging. Six hours after injection, NSCs were tracked. Signal indicating the presence of NSCs was observed in all animals that had received an intraspinal injection of NSCs (Figure 2A-D) with varying flux (Figure 2E). The detectable amount of signal was low at 6 hours, most likely due to factors such as the number of cells injected (100,000 cells), depth of penetration, and presence of multiple tissue layers (fur, skin, fat tissue and muscle). Twenty-four hours after injection, there was minimal signal detected and by day 7 post-injection, no signal was detected in any of the mice. The total flux detected by bioluminescence did not correlate with the number of surviving NSCs subsequently observed in the sectioned spinal cord.

**Figure 2.**
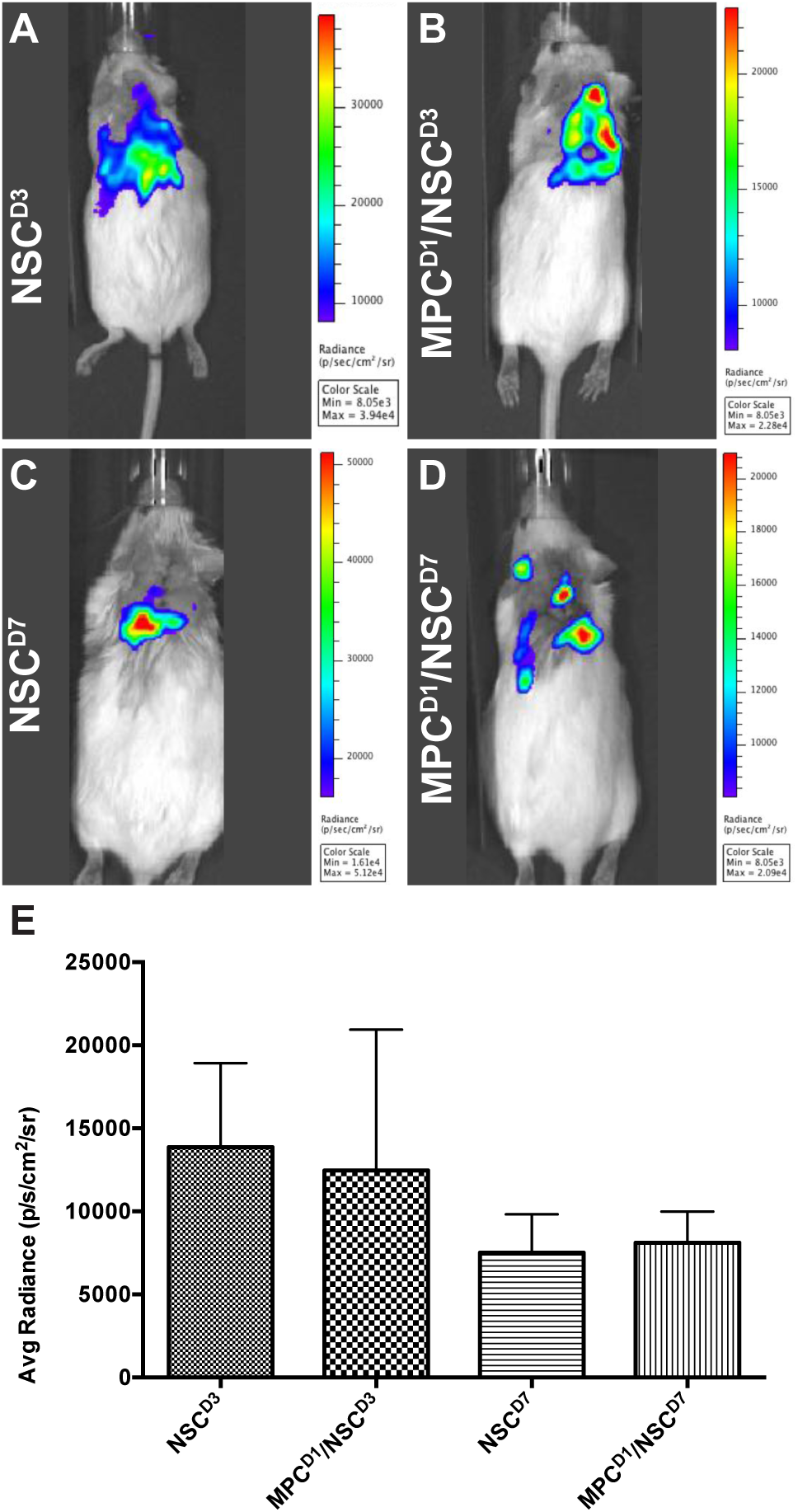
Bioluminescence imaging to track luc-GFP+ NSCs injected after cervical contusion spinal cord injury. Bioluminescence imaging to track luc-GFP+ NSCs injected after cervical contusion spinal cord injury. Imaging at 6 hours post-injection in all mice receiving NSCs. Irrespective of whether MPCs were injected before NSC transplantation, a bioluminescence signal was detected around the injection site in the mice (A, B, C, D) with varying signal dispersion. (E) Bioluminescence emissions was measured and averaged per group. There was no statistically significant difference detected between the groups.

### Dual-IV and IS injection of MPCs and NSCs in unilateral SCI shows limited capacity to improve forelimb function

The mouse cylinder test was used to gauge changes in behavioral outcome as a result of the treatment. There were no significant differences observed in behavioral changes in any of the treatment groups when compared to their controls (Figure 3A, B). Right paw usage varied between 25-35% in all the groups.

**Figure 3.**
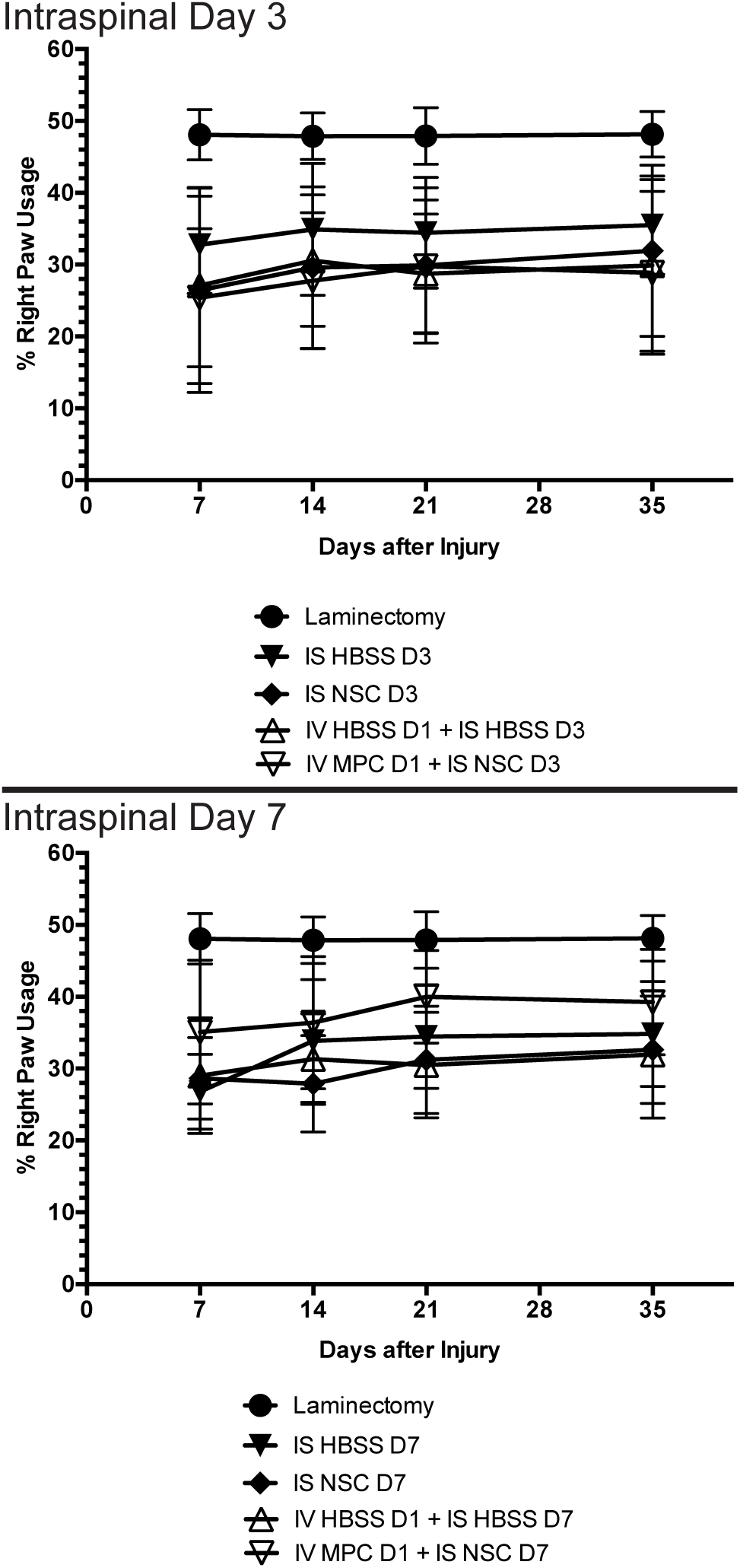
Mean right paw usage over 6 weeks after NSC transplantation in cervical spinal cord injury. Mouse cylinder test results for (A) mice with contusion injury receiving NSCs at D3 and (B) mice with contusion injury receiving NSCs at D7. No statistically significant difference was observed at any time point, irrespective of whether an intravenous injection of MPCs was administered beforehand.

### Intraspinal injection of NSCs results in microcyst formation at the injection site

Histological analysis of the spinal cord tissue showed that micro-cysts formed after intraspinal injection of NSCs (for examples see Figure 5A, 5C, 7B, 8B, 8C). This was not observed when mice were only IV injected with MPCs [18]. The micro-cysts may have formed due to an intraspinal injection artifact caused by the insertion of the needle into the spinal cord. This micro-cyst created a defect in the cord that was devoid of any cells or scar, regardless as to whether NSCs or HBSS were injected.

### GFP+-NSCs isolated from the sub ventricular zone can survive and differentiate in the cervical injured spinal cord

Six weeks after the initial injury, mice were sacrificed, and histological analyses were carried out on the spinal cord tissue. Detection with anti-GFP antibody showed that some of the injected NSCs had survived and differentiated in the host spinal cord (see Figure 5, 6, 7, 8). The distribution of GFP+ NSC cells counted in each section is shown in Figure 4A. There was a statistically significant difference observed in the number of GFP+ NSCs in NSCD3 compared to NSCD7 (p=0.008), NSCD7 compared to MPCD1/NSCD7 (p<0.001) and MPCD1/NSCD3 compared to MPCD1/NSCD7 (p=0.0011). However, in all cases, the number of surviving GFP+ NSCs was smaller to the number of cells that were initially injected. In eight sections counted per animal, the highest cell count in any one section was 63 GFP+ NSCs.

**Figure 4.**
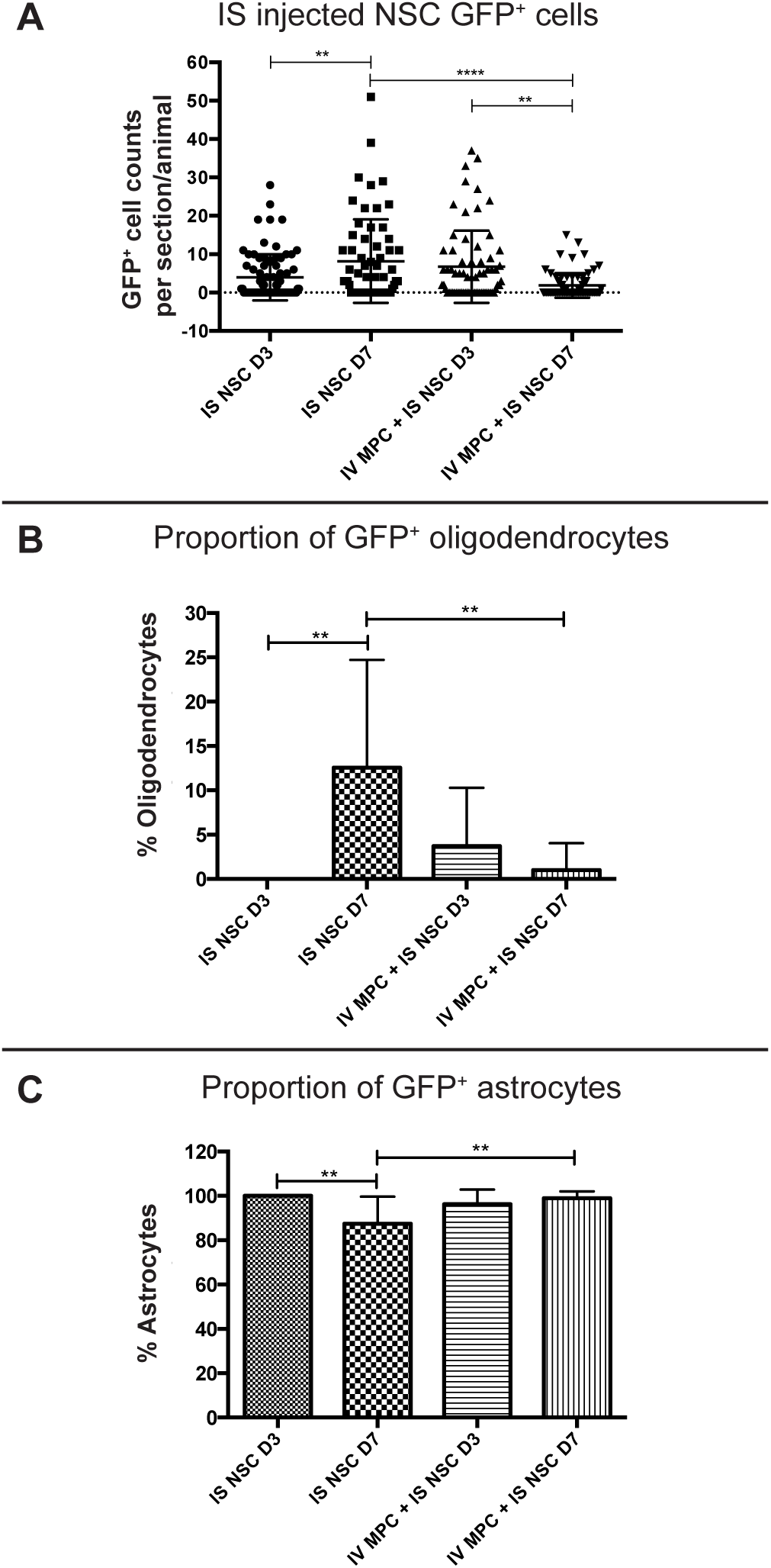
GFP+ NSC cell counts in the spinal cord 6 weeks after injury. (A) The number of GFP+ NSCs in the processed spinal cord tissue was counted per section, per mouse, and displayed as a dot plot summary. (B) The number of GFP+ NSCs that differentiated into oligodendrocytes was counted and plotted as a percentage from the total cells. (C) The number of GFP+ NSCs that differentiated into astrocytes was counted and plotted as a percentage from the total cells. The percentage of oligodendrocytes and astrocytes are inversely proportional as no transplanted NSCs differentiated into neurons.

GFP+ NSCs were labeled with anti-APC, anti-GFAP and anti-βIII tubulin marker labeling for oligodendrocytes, astrocytes and neurons respectively. The GFP+ NSCs were identified as either astrocytes or oligodendrocytes. There was no evidence of any of the GFP+ NSCs differentiating into βIII tubulin+ neurons. In the NSCD3 group, there was no evidence of any GFP+ NSCs differentiating into oligodendrocytes (Figure 4B). The group with the most oligodendrocytes observed was the NSCD7 group. There was a statistically significant difference in the percentage of oligodendrocytes observed between the NSCD7 and MPCD1/NSCD7 (p=0.0081) groups, and a statistically significant difference in the percentage of oligodendrocytes observed between NSCD3 and NSCD7 (p=0.0037). Overall, the majority of the intraspinal injected NSCs differentiated into GFAP+ astrocytes (Figure 4C).

### Intraspinal injection at day 7 increased survival of NSCs and promoted oligodendroglia differentiation

Among the four treatment groups, NSCD7 provided the best integration and survival of injected NSCs (mean = 46.625 cells/mouse), with 12.55% of the cells differentiating into oligodendrocytes (Figure 4, 5). Differentiated astrocytes integrated around the lesion and injection site and were also in close contact with other GFP+ astrocytes (Figure 5Aiv, Biv, Civ). Differentiated oligodendrocytes migrated to the periphery of the spinal cord, extended processes, and were distributed evenly from other GFP+ oligodendrocytes (Figure 5Avii, Bvii). GFP+ oligodendrocytes were co-labeled with anti-APC (Figure 5D).

**Figure 5.**
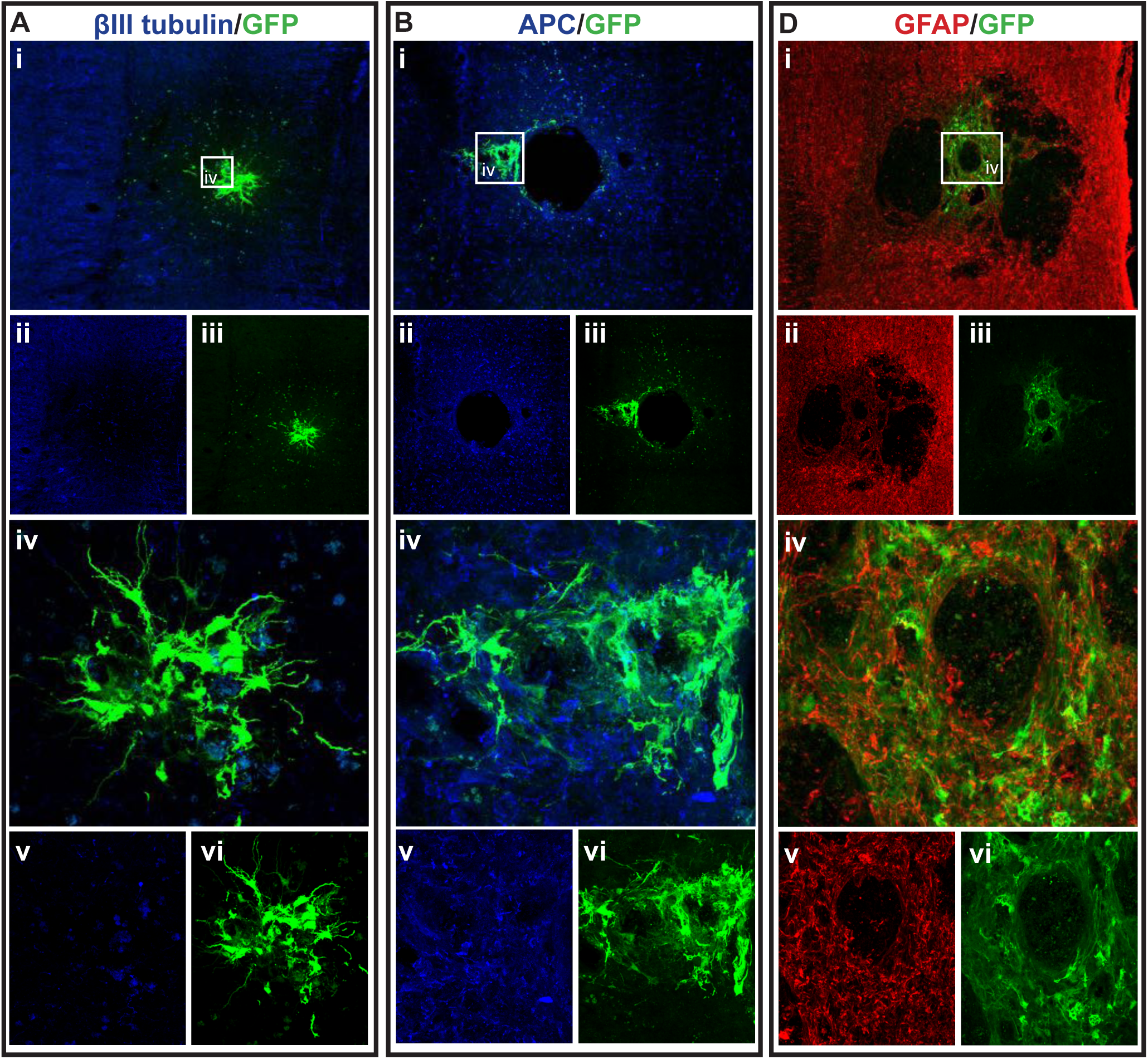
GFP+ NSCs in spinal cord sections of NSCD3 mice. Gross overview of surviving GFP+ NSCs transplanted at day 3 after cervical spinal cord injury showing (Ai, ii, iii) βIII-tubulin and GFP, (Bi, ii, iii) APC and GFP and (Ci, ii, iii) GFAP and GFP. High magnification images showing differentiated (Aiv, v, vi) GFP+ astrocytes with βIII-tubulin, (Biv, v, vi) GFP+ astrocytes and APC and (Civ, v, vi) GFP+ astrocytes and GFAP. Scale bars for images (i, ii, iii) are 200 µm and all other images are 50 µm.iii

### Injection of IV MPCs increased NSC differentiation into oligodendrocytes in mice at Day 3 injection timepoint

Integration and viability of intraspinal injected NSCs at D3 was similar, regardless of whether IV injection of MPCs was carried out at D1 (Figure 4A). There was a mean of 31 cells/section in the NSCD3 group versus a mean of 35 cells/mouse in the MPCD1/NSCD3 group. However, NSC differentiation changed with intravenous injection of MPCs at D1 across the Day 3 and Day 7 time points. For instance, no oligodendrocyte differentiation was observed in the NSCD3 group (Figure 4B), whereas in the MPCD1/NSCD3 group, 3.71% of the surviving cells differentiated into APC+ oligodendrocytes (Figure 4B, 6A, 6B, 6C). The cellular pattern of differentiation in MPCD1/NSCD3 group was similar to that observed in the NSCD7 group as GFP+ astrocytes integrated around the lesion and injection site and were in close contact with other GFP+ astrocytes (Figure 6Aiv, Div), while differentiated oligodendrocytes migrated to the peripheral white matter of the spinal cord and extended processes (Figure 6Avii, BiV, D). The cellular pattern of astrocytic differentiation in the NSCD3 group was like the NSCD7 and MPCD1/NSCD3 groups, with the GFP+ astrocytes integrated around the lesion (Figure 7iv) and injection site (Figure 7Biv) and in close contact with other GFP+ astrocytes (Figure 7).

**Figure 6.**
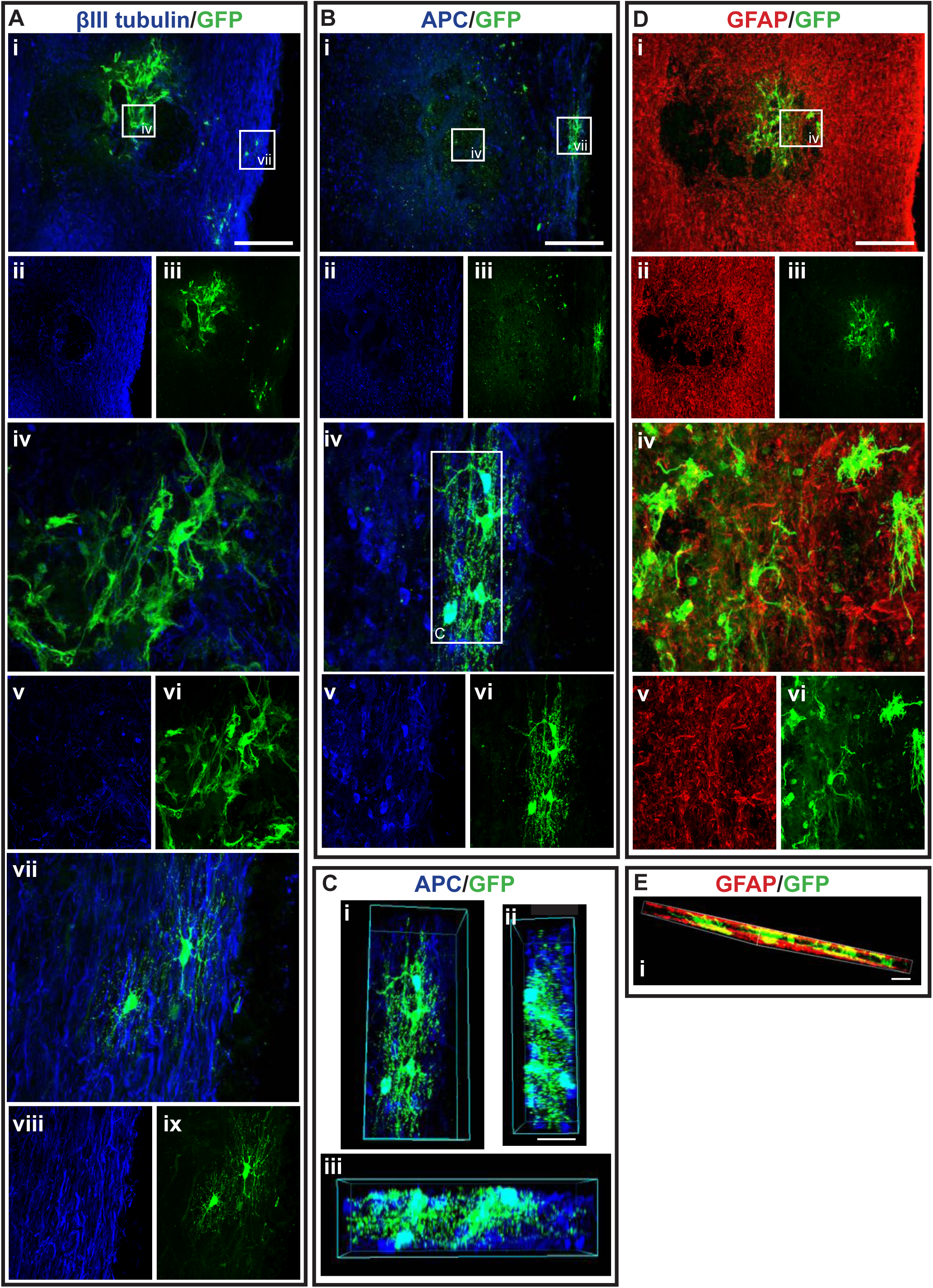
GFP+ NSCs in spinal cord sections of MPCD1/NSCD3 mice. Gross overview of surviving GFP+ NSCs transplanted at day 3 after cervical spinal cord injury and intravenous injection of MPC at D1 showing (Ai, ii, iii) βIII-tubulin and GFP, (Bi, ii, iii) APC and GFP and (Ci, ii, iii) GFAP and GFP. High magnification images showing differentiated (Aiv, v, vi) GFP+ astrocytes with βIII-tubulin, (Avii, viii, ix) GFP+ oligodendrocytes and βIII-tubulin, (Biv, v, vi) GFP+ oligodendrocyte and APC, and (Civ, v, vi) GFP+ astrocytes and GFAP. (Di, ii, iii) High magnification 3D view of GFP+ oligodendrocyte as outlined in white box in (Bvi) from various angles showing co-labeling with APC. High magnification 3D view of (Ei) GFP+ astrocytes and GFAP as outlined in white box in (Ci) and (Eii) 2D view of the same cells with orange cross showing the cross-section of cells observed on side. Scale bars for images (i, ii, iii) are 200 µm and all other images are 50 µm.

**Figure 7.**
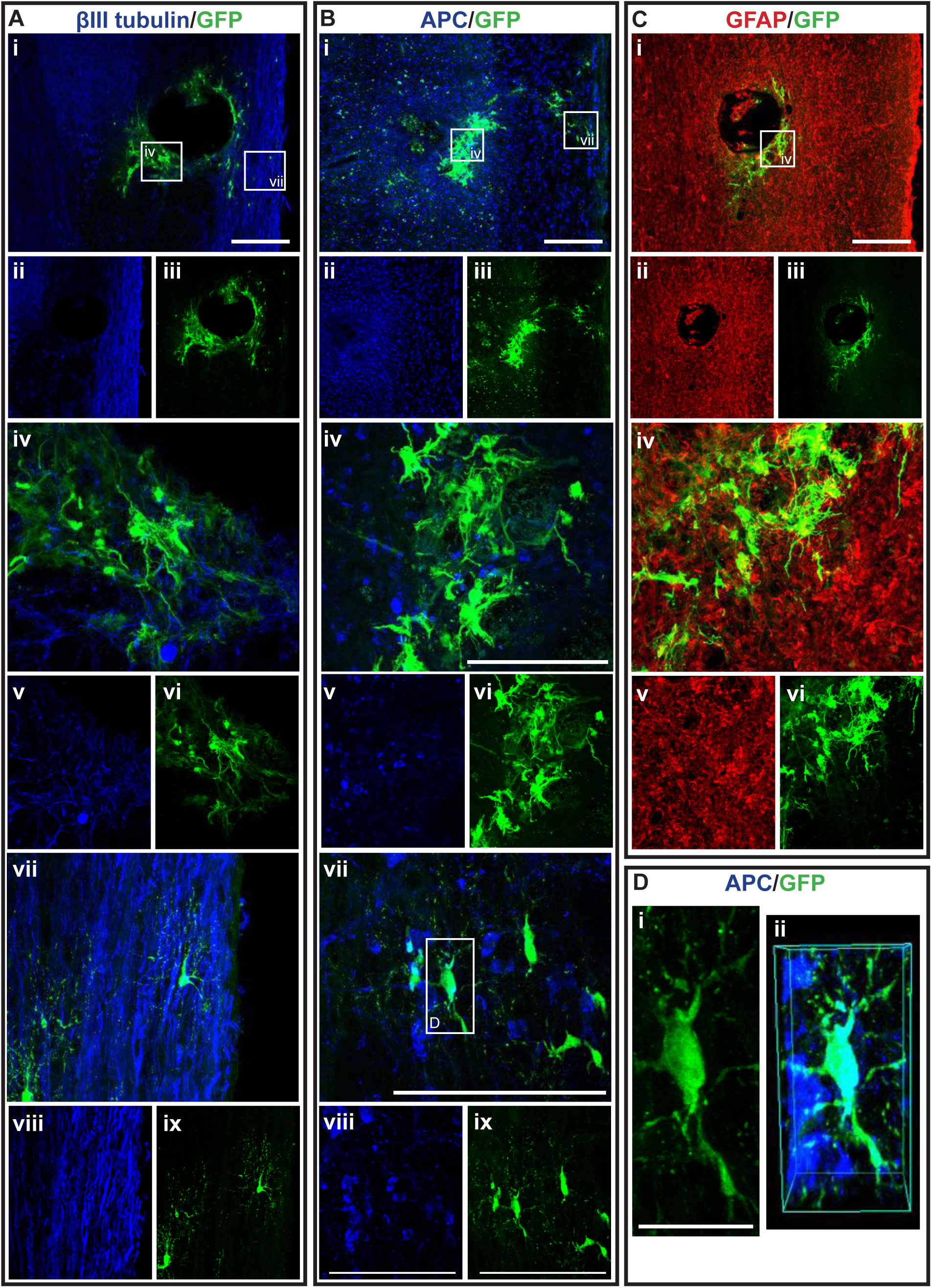
GFP+ NSCs in spinal cord sections of NSCD7 mice. Gross overview of surviving GFP+ NSCs transplanted at day 7 after cervical spinal cord injury showing (Ai, ii, iii) βIII-tubulin and GFP, (Bi, ii, iii) APC and GFP and (Ci, ii, iii) GFAP and GFP. High magnification images showing differentiated (Aiv, v, vi) GFP+ astrocytes with βIII-tubulin, (Avii, viii, ix) GFP+ oligodendrocytes and βIII-tubulin, (Biv, v, vi) GFP+ astrocytes and APC, (Bvii, viii, ix) GFP+ oligodendrocytes and APC and (Civ, v, vi) GFP+ astrocytes and GFAP. High magnification view of (Di) GFP+ oligodendrocyte as outline in white box in (Bvii) and (Dii) shows the 3D view of the same oligodendrocyte co-labeled with APC. Scale bars for images (i, ii, iii) are 200 µm and all other images are 50 µm, except for (Di) where the scale bar was 10 µm.

### Combination of IV injection of MPCs with intraspinal injection of NSCs at D7 resulted in reduced NSC survival and differentiation capacity

MPCD1/NSCD7 group showed the least successful NSC survival integration when compared to all the groups, with an average of 8 cells per mouse (Figure 4A). Approximately 1.01% of NSCs in the MPCD1/NSCD7 group differentiated into oligodendrocytes, which equates to around one cell in all the sections counted (Figure 8Bvii). The GFP+ astrocytes in the MPCD1/NSCD7 group were like the other groups, with integration around the lesion and injection site, while being in close proximity to other GFP+ astrocytes (Figure 8A, C). However, GFP+ oligodendrocytes had migrated caudally from the injection site, rather than to the periphery of the spinal cord (Figure 8B).

**Figure 8.**
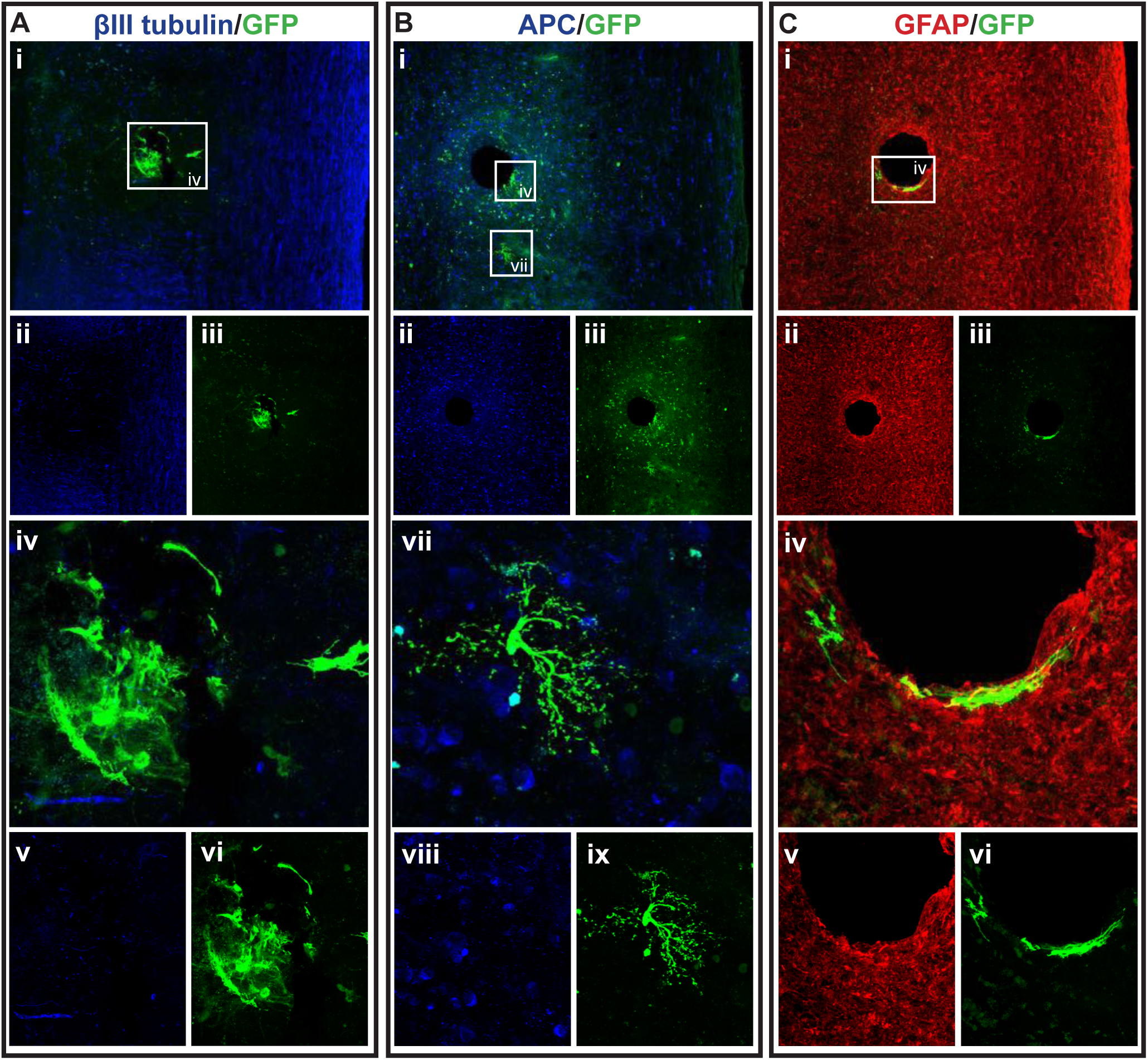
GFP+ NSCs in spinal cord sections of MPCD1/NSCD7 mice. Gross overview of surviving GFP+ NSCs transplanted at day 7 after cervical spinal cord injury and intravenous injection of MPC at D1 showing (Ai, ii, iii) βIII-tubulin and GFP, (Bi, ii, iii) APC and GFP and (Ci, ii, iii) GFAP and GFP. High magnification images showing differentiated (Aiv, v, vi) GFP+ astrocytes with βIII-tubulin, (Biv, v, vi) GFP+ oligodendrocyte and APC and (Civ, v, vi) GFP+ astrocytes and GFAP. Scale bars for images (i, ii, iii) are 200 µm and all other images are 50 µm.

### Distribution of Proteoglycan deposition and Schwann cell ingression

Deposition of proteoglycan and invasion of Schwann cells was assessed using CS-56 and p75, respectively. In the NSCD3 group, the degenerating lesion epicenter showed both the presence of CS56 proteoglycan staining (Figure 9A), but also p75+ Schwann cells (Figure 9B), that had distributed into the lesion. The staining for CS-56 and p75 was overlapping at the injury site, suggesting Schwann cells expressed proteoglycans after injury to the spinal cord (see arrows). Co-expression of Schwann cells and CS-56 has been previously described when Schwann cells were transplanted into a transected spinal cord [27]. In the MPCD1/NSCD3 group, there was an increase in expression of CS-56 and p75 co-expression surrounding the core of the lesion (Figures 9D, E and F, see circle outline). The numbers of Schwann cells were larger in this group but did not enter the lesion core parenchyma, unlike in the NSCD3 group alone. NSCD7 transplants showed a larger degeneration of tissue in the lesion site (*). Schwann cells expressing p75 and CS-56 was observed in the lesion (Figure 9G, H and I). Noticeably, when MPCD1 were combined with NSCD7 intraspinal injections the size of the degenerating lesion core was smaller but did not change the dual expression of p75+ Schwann cells with CS-56 in the center of the lesion (see Figure 9J, K and L).

**Figure 9.**
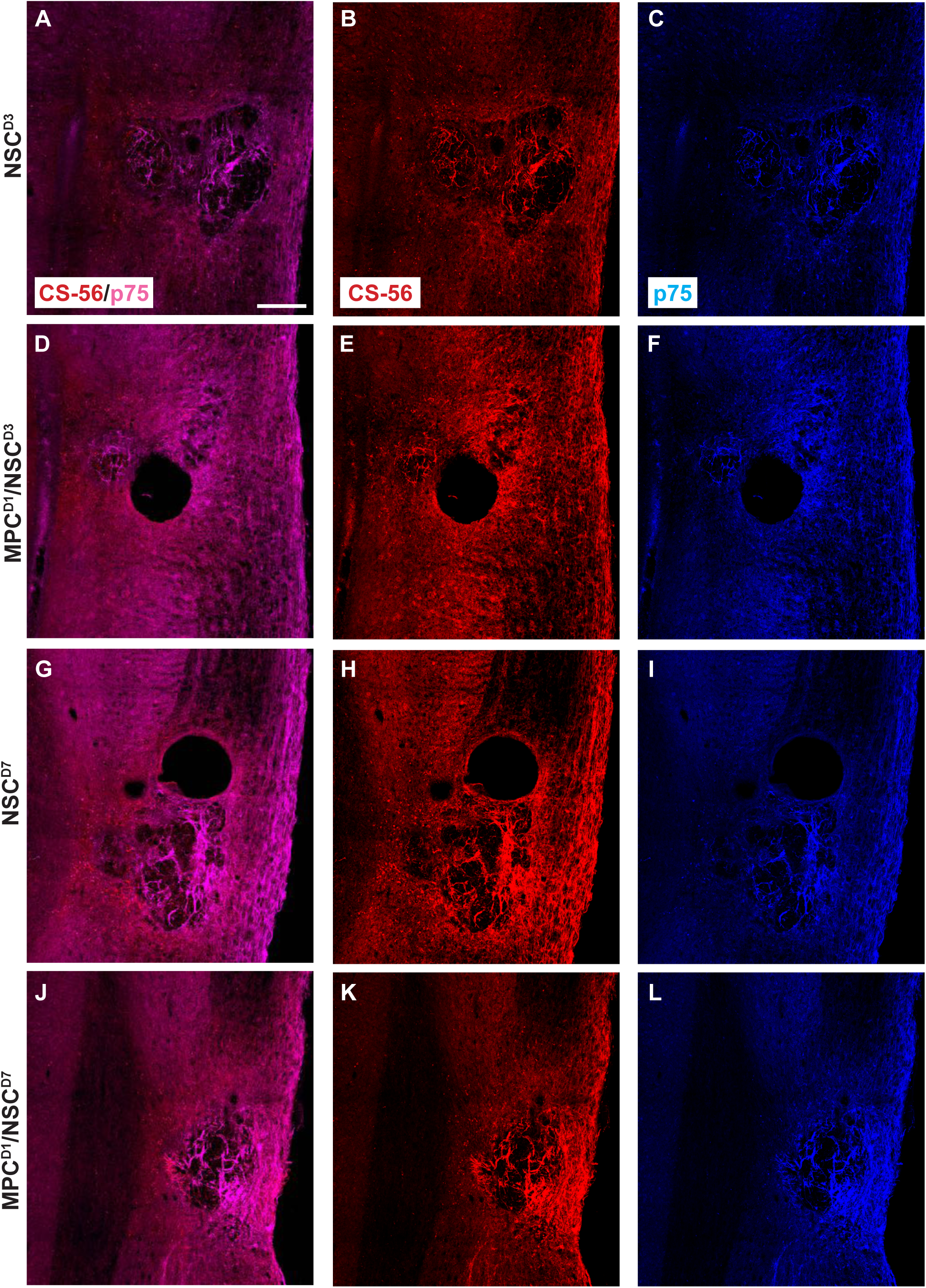
Chondroitin sulfate proteoglycan and p75 profiles in cervical contusion injury treatment groups 6 weeks after injury. Gross overview of chondroitin sulfate proteoglycan deposit stained with CS-56 and Schwann cell infiltration identified by anti-p75. In NSCD3 group, dual label of CS-56 and p75 is observed in (A), while (B) and (C) shows the respective single channel image. White arrow shows examples of area with dual label of CS-56 and p75. In MPCD1/NSCD3 group, dual label of CS-56 and p75 is observed in (D), while (E) and (F) shows the respective single channel image. White dashed circle shows area with dual label of CS-56 and p75. In NSCD7 group, dual label of CS-56 and p75 is observed in (G), while (H) and (I) shows the respective single channel image, * shows the lesion site. In MPCD1/NSCD7 group, dual label of CS-56 and p75 is observed in (J), while (K) and (L) shows the respective single channel image. Scale bar = 200 µm.

### Extracellular matrix, deposition and pericyte infiltration at the site of NSC transplantation is limited by prior injection of MPCs at Day 1

Vasculogenesis and myelin integrity was assessed in all groups using tomato lectin (vessels) and fluoromyelin (myelin). Following supraspinal injections of NSCD3, tissue sections showed an intense halo of new vessels at the lesion site and surrounding the injection site (*) (Figure 10A). Noticeably the core lesion site (arrows) was filled with dense tomato lectin+ structures. Fluoromyelin staining of the same sections showed a demyelinated core (*) surrounded by a demyelinated zone emanating from it (see arrows) (Figure 10B). Neighboring sections stained for the pericyte marker PDGFrβ showed an intense deposition in the lesion/injection site, indicating dissociation from intact blood vessels and formulating scar filling tissue (Figure 10C). Staining in the MPCD1/NSCD3 group indicated similar new blood vessels stained with tomato lectin but the extent of demyelination, even accounting for the injection artifact, was less (*) (Figures 10D and E). PDGFrβ immunostaining showed pericytes filling the lesion area (Figure 10F). However, in the NSCD3 group the PDGFrβ immunostaining showed a fascicle-like morphology (Figure 10C, arrow) but in the MPCD1/NSCD3 group, the PDGFrβ immunostaining is very densely concentrated with no distinct morphology (Figure 10F, arrow).

**Figure 10.**
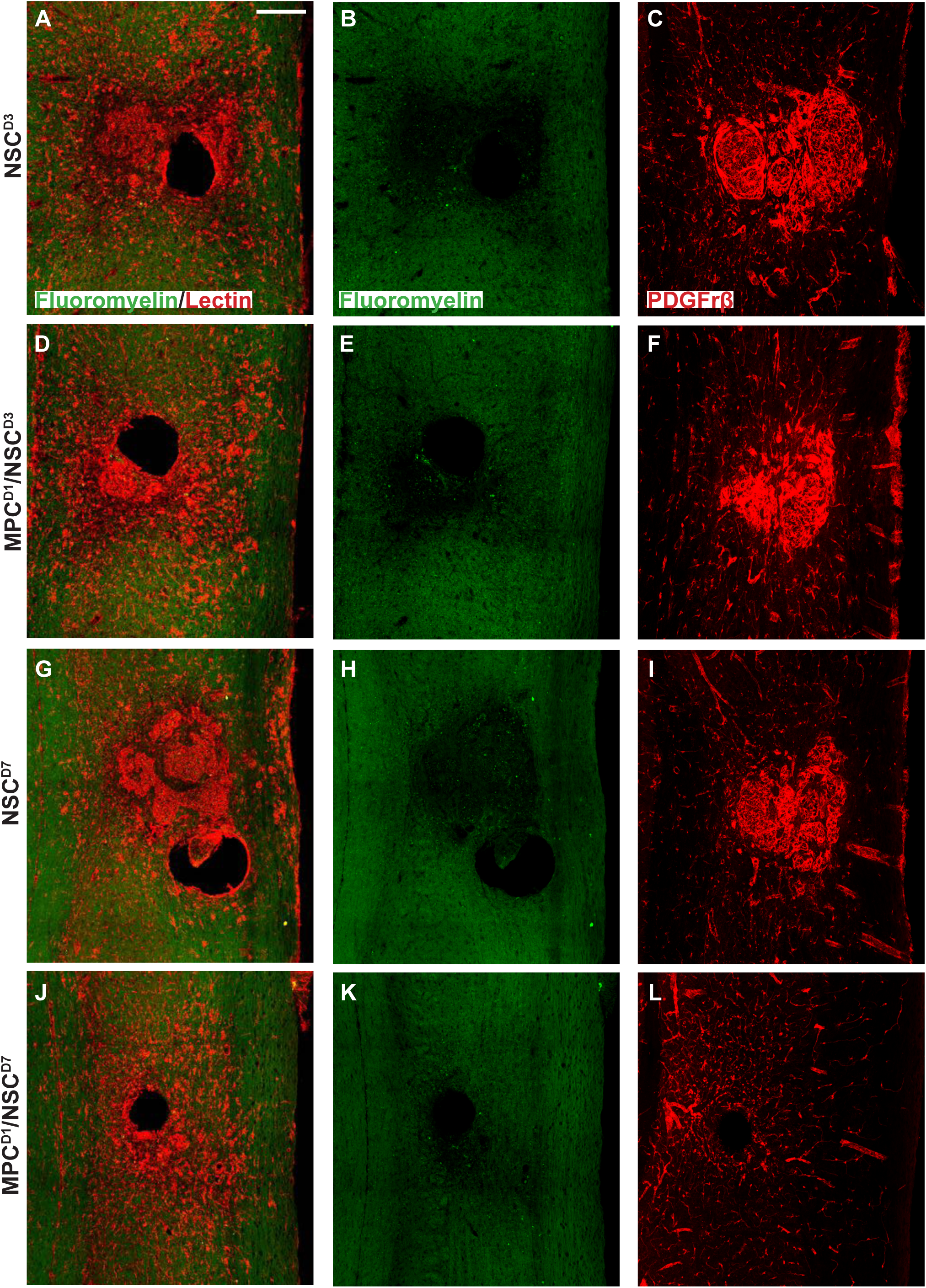
Vascularization profiles in cervical injury treatment groups 6 weeks after injury. Gross overview of demyelination profile, blood vessel formation and pericyte expression around the injection and injury site as shown by fluoromyelin, tomato lectin and PDGFrβ labeling respectively. (A) shows fluoromyelin and tomato lectin co-label with arrows denoting the injury site and * showing the injection site in NSCD3 group while (B) shows the same section with only fluoromyelin expression with arrows showing the edge of the injury site and extent of demyelination and * showing the demyelinated core. In (C), PDGFrβ expression is shown extensively around the injury site with a swirly morphology in the NSCD3 group. (D) shows fluoromyelin and tomato lectin expression in MPCD1/NSCD3 group while (E) shows the same section with only fluoromyelin expression with * showing the demyelinated region. (F) shows the PDGFrβ expression around the injury site in the MPCD1/NSCD3 group with arrow showing dense region of PDGFrβ expression. (G) shows fluoromyelin and tomato lection expression in NSCD7 groups while (H) shows the same section with only fluoromyelin expression. (I) shows PDGFrβ expression around the injury site in the NSCD7 group. (J) shows fluoromyelin and tomato lectin expression in MPCD1/NSCD7 group while (K) shows the same section with only fluoromyelin expression. (L) shows PDGFrβ expression around the injury site in the MPCD1/NSCD7 group. Scale bar = 200 µm.

NSCD7 tissue showed the presence of concentrated vessel networks in the demyelinated zone of the injury site labeled by tomato lectin (Figure 10G). The vessels covered a large grey matter area at the lesion site. The extent of demyelination around the lesion site corresponded to the areas of most concentrated vessel networks (Figure 10G and H). Figure 10I shows PDGFrβ+ pericytes filling the demyelinated zone, indicating that pericytes within the lesion site correspond to new blood vessel formation/scar. In the MPCD1/NSCD7 group, there was a lack of intensively stained tomato lectin+ structures previously observed in the other 3 groups (Figure 10 J, arrows). Fluoromyelin staining also indicated a reduction in myelin loss (Figure 10K). Immunostaining using PDGFrβ for pericytes (Figure 10L) in an adjoining section showed very few PDGFrβ+ pericytes within the lesion and injection site. The PDGFrβ coverage in white and grey matter surrounding the injury zone appeared as intact vessels.

### In vitro MPC/NSC co-cultures shows the increased Oligodendrocyte cell fate for NSCs

Isolated NSCs and MPCs were co-cultured by non-contact culture inserts (see Supp Fig 1) and fed for 0h, 6h, 12h, 1d, 3d, 7d and 10days. Immunocytochemistry was used after cell fixation to show proportion of NSCs differentiated to glial and neuronal phenotypes including GFAP, Beta-III Tubulin (TUJ1), Map2, O4, NG2 and APC at the time points above. Astrocyte (GFAP+) number is not statistically (p<0.05) different at day 10 with or without the presence of MPCs (Supp Fig 1A). Tuj1+ neurons was also not significant at day 10 when compared to NSC only cultures (supp Fig 1B). Map2 staining also showed no increase in the number of MAP2+ cells derived from NSCs in the presence of MPCs (supp Fig 1C). O4+ profiles were statistically increased from 6h through day 10 when compared to NSC only cultures (supp Fig 1D). The same statistical increase in NG2+ cells were also seen in co-cultures with MPCs increasing from 6h through to 10D (supp Fig 1E). Notably, at day 7, quantification of mature oligodendrocyte APC marker shows a large statistical increase in APC+ oligodendrocytes in vitro at day 7 when NSCs are exposed to the MPC secretome (supp Fig 1F). Without the presence of MPCs on the culture inserts, the NSCs did not differentiate into oligodendrocytes but rather went to default astrocytic (GFAP+) and neuronal (TUJ1+) cell fate. This suggests that the MPCs are expressing secretory factors that drive the oligodendrocyte phenotype. Further analysis of MPC secretome is underway to ascertain key candidates.

## 4. Discussion

This study examined a dual transplantation strategy using intravenous mesenchymal progenitor cells (MPCs) and intraspinal NSCs to therapeutically treat adult mice following a cervical model of SCI. Luciferase/GFP+ NSCs (Luc/GFP+) were delivered intraspinal at the C5 level of the spinal cord following a contusion injury and monitored using real time BLI followed by immunofluorescence histology. This live and fixed cell tracking enabled us to ascertain the potential of the Luc/GFP+ NSCs to differentiate to glia or neurons. In addition to the intraspinal NSC transplants animals we also included pre-conditioned mice with intravenous MPCs at 24 hours following SCI or not. Preconditioning mice with MPCs [19] was to provide a reduced inflammatory environment for increased Luc/GFP+ NSC survival and influence NSC differentiation into oligodendrocytes, astrocytes or neurons. The differential outcomes observed between the Day 3 and Day 7 Luc/GFP+ NSC transplantation points towards a critical timing element in SCI intervention strategies. The enhanced integration of NSCs at Day 3, as opposed to Day 7 post-injury, suggests that the early post-injury environment, influenced by MPC treatment, is more conducive to NSC integration and survival. It highlights the role of early immune signaling in determining the success of cellular interventions.

### NSC differentiation into oligodendroglia increased in the presence of MPCs

Increased oligodendrocyte differentiation at Day 3 transplantation, when in the presence of IV-delivered MPCs, indicated a pro-oligodendrocyte secretome. This observation aligns with our in vitro findings, where MPCs co-cultured with NSCs showed an increased oligodendrocyte cell fate, suggesting that MPCs secrete factors promoting oligodendrocyte differentiation. This aspect opens new avenues for exploring the secretome of MPCs in the context of SCI by enhancing axonal remyelination and improve function. This could be via long distance signaling through exosome secretion by MPCs [28, 29].

Although we did not directly investigate this possible mechanism, our MPCs lodged in the lungs post-transplant still lead to an increased oligodendrocyte lineage in transplanted NPCs suggesting that exosomal signaling could be a possibility in the increase of oligodendrocyte lineage not just a direct secretion of factors.

### NSCs isolated from the sub ventricular zone can survive and differentiate in the cervical injured spinal cord

Our study demonstrates the resilience and plasticity of NSCs isolated from the subventricular zone, as these cells were able to survive and differentiate within the injured cervical spinal cord. However, the predominant differentiation is into astrocytes and oligodendrocytes, with an absence of neuronal differentiation, raises questions about the intrinsic and extrinsic factors guiding NSC fate decisions in the SCI environment.

### Intraspinal injection at day 7 increased survival of NSCs and promoted oligodendrocyte differentiation

The enhanced survival and oligodendrocyte differentiation of NSCs observed in the Day 7 intraspinal injection group underscore the significance of the post-injury time window in determining transplantation outcomes. The preferential differentiation into oligodendrocytes suggests a conducive environment for remyelination at this stage. However, the minimal neuronal differentiation points to a limited capacity for direct neuronal circuit reconstruction via NSC transplantation alone. Future investigations should focus on understanding the mechanisms that promote oligodendrocyte differentiation at this stage and how they can be manipulated to support a more diverse range of cell fates, including neurons.

### Injection of IV MPCs altered NSC differentiation in mice receiving intraspinal injection of NSCs at D3

Three days post-injury marks a highly inflammatory period in the injured spinal cord characterized by an influx of microglia and macrophages [22]. Historically, cellular intraspinal transplantation has been carried out one to two weeks post injury when the inflammatory response is less active/reduced [30, 16, 17, 31]. By modulating the inflammatory response with an early IV injection of MPCs, we increased NSCs viability, integration and differentiation when transplanted at D3 post-injury. NSCs differentiated into APC+ oligodendrocytes when MPCs were delivered at D1 prior to intraspinal NSC transplants, something which was not observed in the NSC D3 transplant alone. This differentiation could have been via the MPC secreted factors mentioned above rather than solely a positive adjustment of the immune environment directly.

### Combination of IV injection of MPCs with intraspinal injection of NSCs at D7 resulted in reduced NSC survival and differentiation capacity

We demonstrated in this study that intraspinal transplantation of NSCs at D7 post-injury alone provided the best integration and oligodendrocyte differentiation at a time when microglial activity has peaked. Our results did not support the hypothesis that an additive beneficial effect would occur when two cell types were transplanted: IV injection of MPCs at D1 post-injury reduced the survival, integration and oligodendrocyte differentiation of intraspinal delivered NSCs at D7. The microenvironment of the spinal cord following IV MPCs delivery at Day1 was reduced in suitability for the integration of subsequent intraspinal NSC transplantation at D7 post-injury. From our previous work, IV injection of MPCs at D1 resulted in a smaller injury site, altered glia and vascularization response and the effect occurs rapidly after injection [18]. The reduction in tissue degeneration and vasculogenesis by the treatment of MPCs appears to have reduced NSCs survivability. We also report here that changes in levels of tomato lectin, fluoromyelin, PDGFrβ, CS-56 and p75 expression were observed. It is highly likely that this change at the site of injury created an environment that was less suitable for intraspinal transplantation at day 7 post-injury. Previous research has indicated that cellular interactions between the host and transplanted cells are vital to their integration [32, 33], and indeed there exists strong evidence to show that the host SCI niche is limited in its capacity to provide integration sites or growth signals for transplanted cells [12].

Additional strategies such as biomaterials, matrices and growth factors may greatly assist this integration [34–36]. In addition, the number of transplanted cells can also be crucial in determining the rate of engraftment and differentiation [12]. It has been shown that neural-restricted precursor cells can survive in the intact spinal cord [37] but following spinal cord injury, the microenvironment alters the integration, maturation and migration of transplanted cells [38]. Interestingly, transplantation of hippocampal precursor cells in the retina have shown cells incorporating into damaged areas but not in the intact retinae [39]. By decreasing tissue degeneration and inflammatory signals at the injury site by treatment of IV injected MPCs, we postulated that we created a microenvironment at the injury site that was unsuitable for subsequent intraspinal transplantation of NSCs at D7 to promote regeneration and functional recovery after SCI. In addition, the same number of cells - in this study 100,000 NSCs - may be unsuitable to achieve the best integration or differentiation in the host tissue due to the changes in its microenvironment, compared to when MPCs are not involved. Increasing cell number by pretreating NSCs with anti-apoptotic drugs or providing cell substrates may improve NSC integration. However, larger numbers is not a prerequisite for overall functional recovery.

### Changes in proteoglycan deposition, vascular and pericyte/Schwann cell ingression-role in tissue repair

Extracellular matrix, deposition and pericyte infiltration at the site of NSC transplantation is limited by prior injection of MPCs at 24 hours after injury. The influence of MPCs on the spinal cord extracellular matrix and vascular response post-SCI is a crucial finding of our study. The prior injection of MPCs at 24 hours appears to modulate the extracellular matrix composition and pericyte behavior at the site of NSC transplantation. This modulation could be due to the anti-inflammatory and immunomodulatory properties of MPCs, which might alter the local microenvironment to be less hostile for NSC survival and integration. The reduced pericyte infiltration and altered extracellular matrix deposition in the MPC pre-treated groups suggest a potential mechanism by which MPCs may enhance NSC differentiation.

Understanding the interactions between these cellular therapies and the host microenvironment is essential for optimizing combinatorial strategies in SCI repair. Future research should explore the molecular pathways involved in these interactions and how they can be harnessed to improve the efficacy of cellular therapies for SCI.

### Dual-injection of MPCs and NSCs in unilateral SCI shows limited capacity to improve forelimb function

The findings from our dual-injection approach indicate a limited capacity to enhance forelimb functional recovery in the SCI model. This outcome, despite the observed cellular integration and differentiation, highlights a disconnect between cellular and functional repair mechanisms in SCI. The limited functional improvement could be attributed to the complexity of SCI pathophysiology, where cellular replacement alone may not suffice to restore the intricate circuitry and connectivity required for full functional recovery. These results underscore the need to integrate cellular therapies with other strategies, such as neurorehabilitation or bioengineering approaches, to achieve comprehensive functional restoration in SCI.

This study uniquely combined a time-staggered system of two cellular transplantation techniques following mouse cervical SCI: intravenous injection of MPCs at Day 1, and intraspinal injection of NSCs at Day 3 or Day 7. MPCs were used to ameliorate the inflammatory response and prepare a favorable tissue milieu after spinal cord injury [19], in order to encourage NSCs - acting as a bridging or new circuitry substrate - to integrate, survive, differentiate and promote repair of the injured spinal cord following their subsequent transplantation. We demonstrated in this study the following: (1) IV injection of MPCs shifted the transplantation time window for intraspinal NSC transplantation into the spinal cord, (2) IV injection of MPCs improved the long-term viability and integration of NSCs delivered at D3 post-injury, and (3) intraspinal injection of NSCs at D7 was the most successful for NSC integration and oligodendrocyte differentiation in the groups studied.

### “Off the shelf treatment”

This study was designed to examine the effect of prior IV injection of MPCs (24 hours) on the subsequent survival and differentiation of intraspinal Lu/GFP+ NSCs transplantation. Both cell types used in this study were from frozen stocks and were not treated with any growth factors prior to transplantation [19]. In addition, NSCs were injected directly into the injury site without any additional scaffolds. This approach has an added advantage of being a marketable “off the shelf” therapy that would provide more consistent cell phenotypes, translational capacity and reproducibility. No neuronal differentiation was detected at any of the injected time points within our study, with majority of the surviving and integrated NSCs differentiating down an astrocytic lineage. Additionally, our comprehensive in-vitro results support the MPC’s ability to drive towards an oligodendrocyte lineage from NSCs through a secretory mechanism. In accordance with our results, previous studies have shown in vitro induction towards a neuronal lineage and/or modification of the host environment is necessary to facilitate subsequent in vivo differentiation of neurons [40]. In addition, other studies using NSC transplantation into the spinal cord (a homotopic non neurogenic site) have shown that NSCs differentiated into glia, but not neurons [41]; however when transplanted to the hippocampus (a heterotopic neurogenic site) they can generate neurons [42]. Previous data has shown that NSCs have the capacity to differentiate down the neuronal lineage under appropriate niches or conditions, including the use of fetal cells [17], growth factors and scaffolds [35, 43, 44]. Our results show that IV injection of MPCs alone prior to NSC transplantation does not promote robust neuronal integration or neuronal differentiation of NSCs in the host spinal cord. By using two transplantation methodologies that have been individually shown to be of benefit, a more positive outcome was expected to be achieved when combined; most interestingly this proved not to be the case. There was no increase in functional and anatomical improvement. However, this dual transplant methodology did show an increase in NSC integration and differentiation at D3 post-injury, a period of active inflammatory activity in the injured spinal cord. This transplant timepoint is rarely used due to the non-optimal conditions for cell survival. Our study provides another important step in understanding cellular combinatorial approaches using two cell delivery strategies; the timing in which different therapies are administered is also critical to optimize, in order to assure the best possible outcome.

Additional animal studies will be needed to further understand the mechanisms involved in MPC and NSC combinations such as immune cell activation, cell proteomics and tissue niches available for transplanted and endogenous stem cells to survive/integrate into injured spinal cord tissue.

## 5. Conclusions

In conclusion, our study provides valuable insights into the temporal dynamics of cellular responses in SCI and the potential synergistic effects of combining MPC and NSC transplantation. However, the limited functional recovery and challenges associated with cell transplantation emphasize the need for further research. Future studies should focus on optimizing the timing and method of cell delivery, understanding the molecular mechanisms underlying cell interactions, and exploring combinational therapies that can effectively bridge the injured spinal cord and facilitate functional recovery.

**Supplementary Materials:** Figure S1: *in-vitro* differentiation of NSC in MPC secretome.

## Author Contributions

Seok Voon White: Collection and/or assembly of data, Manuscript writing; Yee Hang Ethan Ma: Data Analysis, interpretation, manuscript writing; Christine D Plant: Data analysis and interpretation, Manuscript writing; Alan R Harvey: Data analysis and interpretation, Manuscript writing; Giles W Plant: Conception and design, financial support, Provision of study material, Data analysis and interpretation, Manuscript writing, Final approval of manuscript.

## Institutional Review Board Statement

The animal study protocol was approved by the Institutional Review Board (or Ethics Committee) of Stanford University. APLAC: 26808 (08/13/2020), APB: 2439 (08/21/2021), SCRO: 371 (03/09/2021).

## Acknowledgments

The authors would like to thank Professor Joseph Wu, Stanford University, for providing the GFP-luciferase transgenic mice used in this study. The authors would like to thank the Small Animal Imaging Facility Stanford Center for Innovation in In-Vivo Imaging (SCI3), Stanford University, for use of IVIS Spectrum. We would also like to thank the Behavioral and Functional Analysis Laboratory, Stanford University, for use of their surgical suite and mouse cylinder test set-up. We thank Chris Czisch, department of Neurosurgery, Stanford University, for the technical assistance in completing the project. We thank Cyrus Ho Hin Chan, The Ohio State University, for assisting in generating statistics for *in-vitro* experiment.

## Appendix

**Appendix A1.** Abbreviations and number of animals per group analyzed.

| Group | Abbreviations | n |
| --- | --- | --- |
| Intraspinal HBSS D3 | HBSS<sup>D3</sup> | 10 |
| Intraspinal NSC D3 | NSC<sup>D3</sup> | 8 |
| Intravenous HBSS D1 with intraspinal HBSS D3 | HBSS<sup>D1</sup>/HBSS<sup>D3</sup> | 9 |
| Intravenous MPC D1 with intraspinal NSC D3 | MPC<sup>D1</sup>/NSC<sup>D3</sup> | 9 |
| Intraspinal HBSS D7 | HBSS<sup>D7</sup> | 9 |
| Intraspinal NSC D7 | NSC<sup>D7</sup> | 8 |
| Intravenous HBSS D1 with intraspinal HBSS D7 | HBSS<sup>D1</sup>/HBSS<sup>D7</sup> | 10 |
| Intravenous MPC D1 with intraspinal NSC D7 | MPC<sup>D1</sup>/NSC<sup>D7</sup> | 9 |

