## Supplemental Figure 1 for "Combined Transplantation of Mesenchymal Progenitor and Neural Stem Cells to repair cervical spinal cord injury"

**A Proportion of GFAP⁺ DAPI**

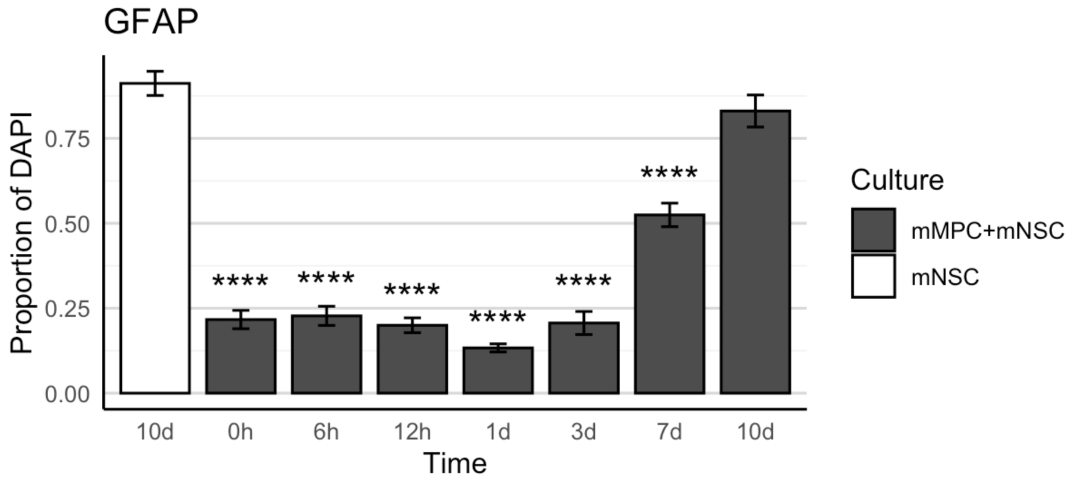


**B Proportion of Tuj1⁺ DAPI**

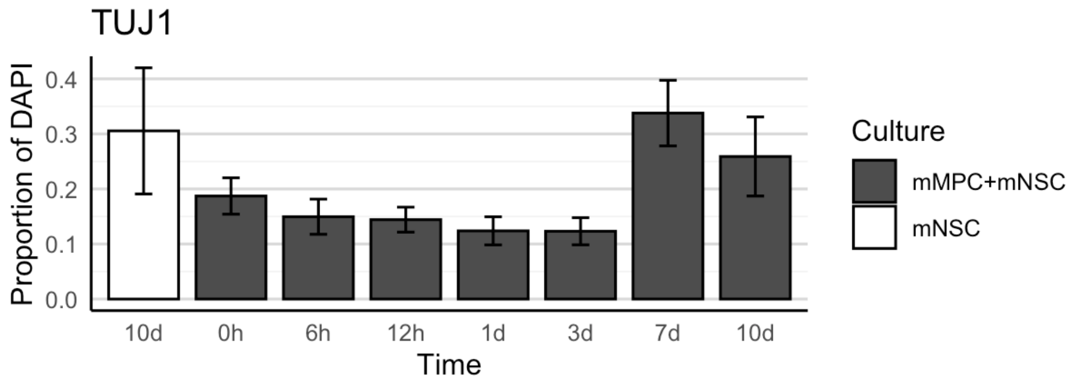

**C Proportion of MAP2+ DAPI**

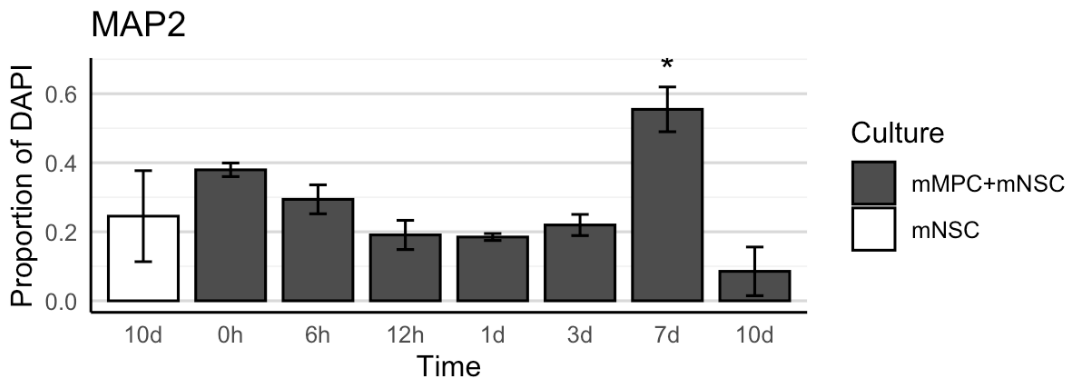


**D Proportion of O4+ DAPI**

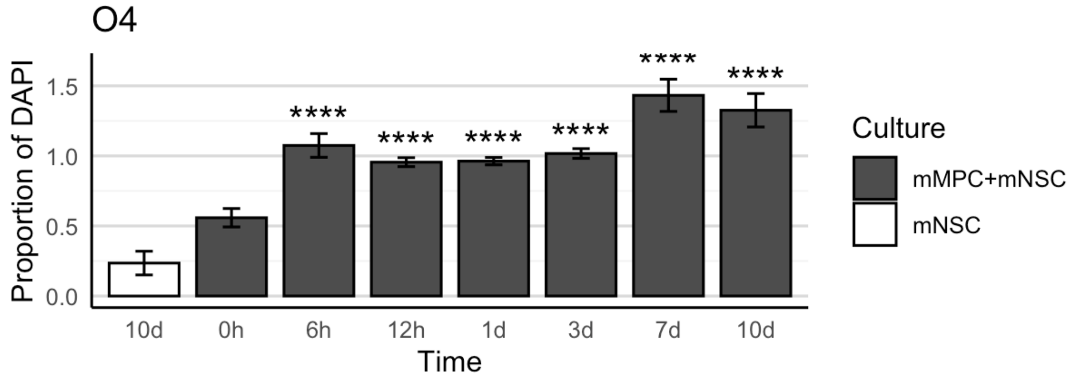

### E Proportion of NG2$^+$ DAPI

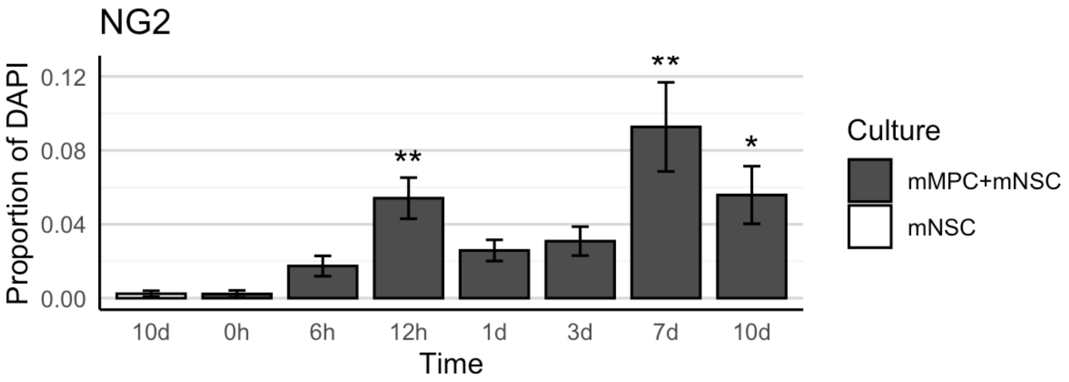


### F Proportion of APC$^+$ DAPI

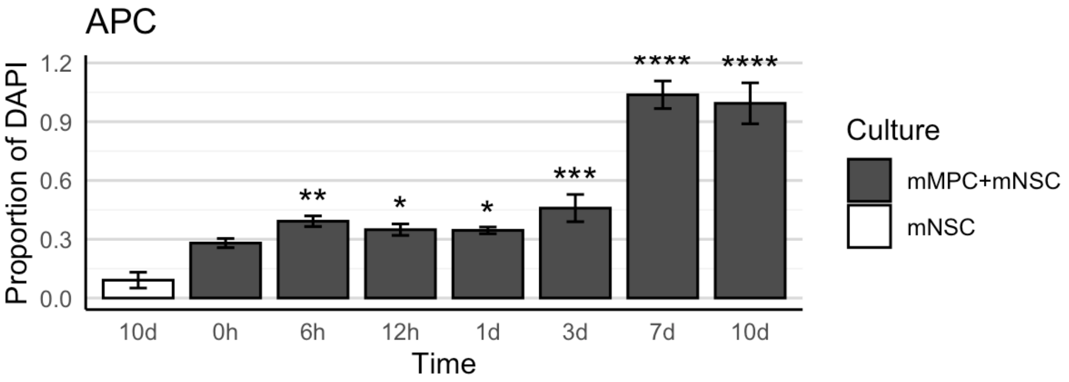


**Supplemental Figure 1. *In-Vitro* differentiation of NSC in MPC secretome**

Proportion of (A) GFAP+ (0h, p= 5.107x 10-15; 6h, p= 5.107x 10-15; 12h, p= 5.107x 10-15; 1d, p= 5.107x 10-15; 3d, p= 5.107x 10-15; 7d, p= 9.681 x 10-14), (B) βIII-Tubulin+ (0h, p= 1.571 x 10-5; 6h, p= 4.319 x 10-6; 12h, p= 3.994 x 10-6; 1d, p= 3.317 x 10-7; 3d, p= 5.540 x 10-7), (C) MAP2+ (7d, p= 1.295 x 10-2), (D) Tuj1+ (0h, p= 1.074 x 10-3; 12h, p= 2.023 x 10-2; 7d, p= 4.167 x 10-2), (E) O4+ (6h, p= 5.816 x 10-10; 12h, p= 3.555 x 10-8; 1d, p= 2.663 x 10-8; 3d, p= 3.029 x 10-9; 7d, p= 9.575 x 10-12; 10d, p= 1.379 x 10-11), (F) NG2+ (12h, p= 5.697 x 10-3; 1d, p= 2.377 x 10-1; 7d, p= 1.800 x 10-3; 10d, p= 1.965 x 10-2), and (G) APC+ (6h, p= 2.595 x 10-3; 12h, p= 3.132 x 10-2; 1d, p= 1.087 x 10-2; 3d, p= 3.945 x 10-4; 7d, p= 0.0000; 10d, p= 0.0000) DAPI+ Nuclei. All groups listed are in mMPC + mNSC cultures, and the statistics are computed in comparison to the 10-day mNSC-only group.
